# Sociality enhances survival in science, especially for female researchers

**DOI:** 10.1101/2020.03.02.973479

**Authors:** Jessica E.M. van der Wal, Rose Thorogood, Nicholas P.C. Horrocks

**Author notes:** The authors declare that no competing interests exist.

## Abstract

Intense competition for limited opportunities means the career path of a scientist is a challenging one, and female scientists in particular are less likely to survive in academia. Collaboration is a key factor in scientific advances, and in social species enhanced sociality improves fitness and longevity. Yet whether sociality influences career progression and survival in science, and how this might differ between genders, is largely unknown. We built authorship social networks from publication records to test how sociality predicts career progression and survival in a cohort of biologists contributing to three international conferences in the 1990s. We show that sociality has the strongest effect for female researchers but, regardless of gender, publishing with many diverse co-authors significantly reduces time to become a principal investigator and increases career duration. Publishing repeatedly with co-authors also enhances career progression in both genders, but reduces career length for men. These findings demonstrate that the value of collaboration extends beyond scientific advances, and can directly benefit the career progression and longevity of research scientists themselves. Efforts to encourage researchers at all levels to invest in collaborations, particularly with female researchers, will help to close the gender gap in science and academia.

## INTRODUCTION

Surviving in academia as a scientist is widely recognised as a challenge, with researchers today experiencing greater job competition, prolonged job insecurity, and declines in career length compared to 50 years ago ^1–7^. For female scientists these challenges are especially acute, since women suffer a variety of inequalities within science compared to men ^8–16^, causing them to leave academia more often, and earlier, than male scientists ^9,17,18^. Lack of diversity – of all kinds – has perverse outcomes, including hampering novel insight and scientific advances ^19,20^. Yet despite much effort to understand the underlying causes (summarised in Fig. 1 of ^21^), disparities between the genders show discouragingly few signs of reducing ^8,9,13,22–25^. Given these obstacles to career attainment and progression, what lessons can we learn from scientists that have survived in science and carved out long careers for themselves?

**Figure 1.**
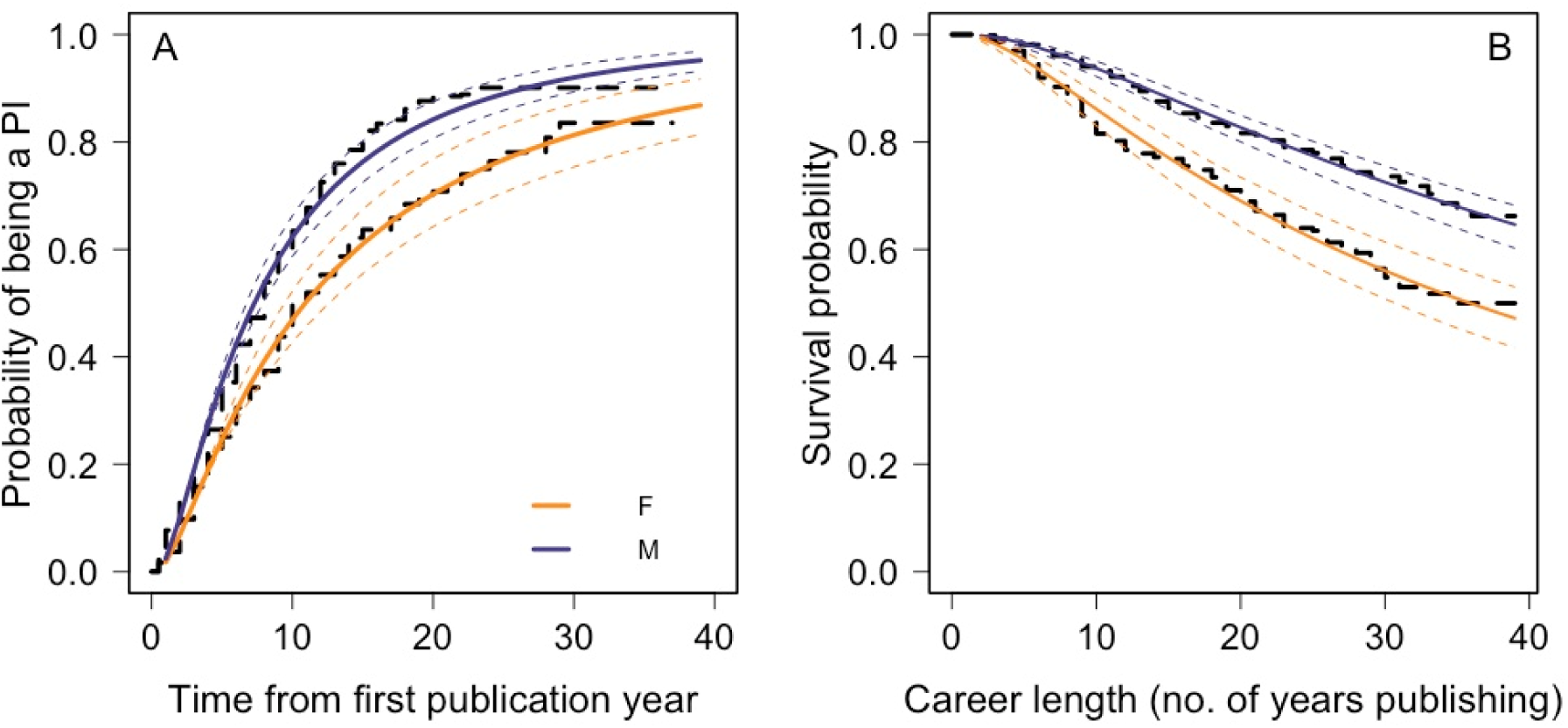
Differences in performance metrics between female (n = 298) and male (n = 637) focal authors, based on their publishing output. Female focal authors in our dataset were (A) less likely and took longer to reach PI status, and (B) had shorter scientific careers than male focal authors, on average (as measured by number of years publishing). Black dashed lines represent Kaplan-Meier survival functions, while the coloured lines represent model predictions (solid lines) with 95% confidence intervals (dotted lines).

One approach to this question is to consider studies that have investigated what underlies longevity and success, and to see if those qualities identified as important also apply to scientists and their careers. For social animals – which scientists surely are ^26–30^ – interacting with peers is a crucial component of lifetime fitness ^31–40^ and lifespan ^41–43^. Several mechanisms have been proposed for how increased sociality could improve fitness and longevity, and that could be relevant to scientists and their careers. These include the role of group level cooperation in acquiring resources, social learning, and the attainment of social status ^37^. In line with these ideas, scientists with larger research networks gain more funding ^44^ and are more highly cited than scientists with fewer co-authors ^29,45–48^, while articles mentioned on social media gain more citations ^49,50^, suggesting that sociality in science also promotes greater access to resources, knowledge of new research, and leads to higher status among scientific peers. In turn, funding and citation rates are both factors that positively influence career progression and longevity in science ^51–54^. The health benefits of sociality are also well-documented, and increased sociality is associated with lower stress levels and reduced incidence of stress-related illness ^41,55–59^. If a larger social network helps researchers to better cope with the stresses associated with establishing and maintaining a career in science and academia ^60–64^, then we might also expect a positive correlation between career lifespan and sociality, with less sociable individuals leaving science earlier.

Here we apply the concepts of studies testing how sociality affects fitness and survival to investigate how social strategies among scientists influence career advancement and duration. Understanding the connections between sociality and career progression and survival, and in particular how this might differ between genders, could shed light on social strategies that researchers may wish to follow, and provide greater understanding of the underlying causes of gender disparities in the sciences. Previous studies have demonstrated the value of research collaborations for academic success (^48,65–70)^, and also that social strategies may differ between the genders ^71–74^. However, the survivability of scientific careers has attracted only limited attention ^1,75,76^, and the links between sociality, gender, and survival in science are completely unexplored. We used social network analyses to quantify the structural properties of egocentric co-authorship networks constructed from the publication records (>53,000 papers) of over 900 gender-identified international scientists, publishing over almost four decades. In contrast to many previous studies, we made no restrictions in terms of career stage ^72,74,77^, journal ^69,78–80^, or institute ^71^, and included both researchers who subsequently left science, and those who are still active ^76^, allowing us to determine directly the effect of sociality on survival in science.

Our cohort consisted of contributors to three consecutive conferences of the International Society for Behavioural Ecology, which includes researchers working on a diverse range of topics in ecology and evolutionary biology. This cohort was ideal for our study for two reasons: i) multi-authored papers are the norm in this field ^69^, which is necessary for the construction of ego-centric networks, and ii) numerous studies of gender biases within ecology and evolutionary biology suggest that, while female researchers remain under-represented in senior or more visible roles ^25,81–85^, biases against female authors in terms of acceptance rates or editorial decisions are not readily apparent ^86–92^ (but see ^25^). Thus, while this field certainly does not display gender equality, for the purposes of our publication-based study, gender bias in publishing is unlikely to be an issue.

Importantly, since female researchers typically publish fewer papers than their male ^71,76,77,79,80,93^, this study, we corrected the metrics we extracted from the social networks we constructed by controlling for variation in the number of papers that these focal authors published, thereby allowing us to account for the productivity gap between the genders. We calculated three measures of author sociality from these publication-corrected social metrics: collaborativeness, which describes the number of unique co-authors a focal author publishes with; consistency, which describes how often, on average, a focal author publishes with the same co-authors; and co-author connectedness, which describes how often the same co-authors appear together across the publications of a focal author. We then used survival analyses (accelerated failure time models; AFT) to directly compare how sociality affects career progression and survival of female and male researchers, a major advancement on previous studies. We expected that scientists who were more social in their approach to publishing – that is, were more collaborative, more consistent, and who had co-authors that were less connected to each other– would i) be more likely to become principal investigators (PIs), ii) become PIs more quickly, and iii) survive for longer in science.

## RESULTS

### Gender differences in publication output-corrected author sociality metrics and in performance measures

Our final dataset included 298 women (32%) and 637 men (68%), and three focal authors for whom gender was unknown (see Supplementary methods S1 for more descriptive statistics of the focal authors). We found that gender significantly affected collaborativeness score (Generalised linear model [GLM], estimate ± SE *=* 0.29 ± 0.07, *t*_933_ = 4.21, *p* < 0.001; Fig. S6D), and males were more collaborative than females (i.e. had a greater number of co-authors on average, given their number of publications). By contrast, gender did not affect consistency (GLM, estimate ± SE = 0.12 ± 0.07, *t*_933_ = 1.73, *p* = 0.08; Fig. S6E) or co-author connectedness (GLM, estimate ± SE = -2.00 ± 1.40, *t*_928_ = -1.43, *p* = 0.15; Fig. S6F). Thus, with the exception of collaborativeness, we found no evidence for underlying gender differences in *how* focal authors in our dataset published their work in terms of their level of sociality. Gender differences *were* apparent, however, in all three measures of academic performance that we investigated. Considering only the effect of gender, male focal authors were significantly more likely to become PIs than their female colleagues (63% of women versus 82% of men; GLM, estimate ± SE = 0.97 ± 0.16, *z*_933_ = 6.12, *p* < 0.001) and did so in about 30% less time (AFT, estimate = 0.69 [0.60, 0.81]). Using the median time of eight years to reach PI status (based on all authors in our dataset that did so), this means that, on average, female focal authors took just over six months longer to reach this career stage than did male focal authors (restricted mean survival time, between-group contrast: M - F = -0.51 years [-0.84, -0.20], *p* = 0.002; Fig. 1B).

Being male also increased the likelihood of still being in science, on average, by 69% (AFT, estimate = 1.69 [1.39, 2.05]). Thus, female focal authors had a mean ± SE career length of 22.16 ± 0.56 years, while for men it was 25.90 ± 0.34 years. Over the maximum 39-year career length of any focal author in our dataset, this equates to a female focal author losing nearly 4.5 years of career time, on average, compared to a male focal author (restricted mean survival time, between-group contrast: M - F = 4.48 years [2.74, 6.23], *p* < 0.001). This is a reduction in career length of 11% compared to male focal authors (Fig. 1A).

### Effect of author sociality on PI status, time to PI, and career length

#### i) Likelihood of becoming a PI

After accounting for gender, all three measures of authorship sociality significantly predicted the likelihood of focal authors achieving PI status. In the case of collaborativeness and consistency, the relationship was positive, such that focal authors who did become PIs tended to have higher scores for these two author sociality metrics (GLM, collaborativeness: estimate ± SE = 1.75 ± 0.15, *z*_932_ = 11.88, *p* < 0.001; consistency: estimate ± SE = 0.39 ± 0.09, *z*_932_ = 4.33, *p* < 0.001). The reverse was true for co-author connectedness: focal authors with a lower connectedness score were more likely to become PIs than those that had higher co-author connectedness scores (GLM, estimate ± SE = -0.53 ± 0.08, *z*_927_ = -6.33, *p* < 0.001).

#### ii) Time to PI

Collaborativeness significantly and positively influenced how long it took focal authors to become PIs, but there was no effect of gender on this relationship (interaction term: AFT, estimate = 1.17 [0.99, 1.38]). For every increase of one in collaborativeness score, the time to PI decreased by 21% (AFT, estimate = 0.79 [0.73, 0.85]; Fig. 2A, B; Fig. 4). Consistency was also a significant predictor of time to PI. Focal authors with higher consistency scores took significantly less time to become PIs, with the effect of consistency being greater for female focal authors than for male focal authors (interaction term: AFT, estimate = 1.34 [1.14, 1.58]; female focal authors estimate = 0.64 [0.55, 0.74]; male focal authors estimate = 0.86 [0.80, 0.93]; Fig. 2C, D; Fig. 4). Focal authors with high co-author connectedness scores took longer to reach PI status than those with lower scores, and this was dependent on gender (interaction term: AFT, estimate = 0.84 [0.72, 0.99]). An increase of one unit in co-author connectedness score decreased the time to PI by approximately 37% in women (AFT, estimate = 1.37 [1.16, 1. 60]; Fig. 2E; Fig. 4) but only 13% in men (AFT, estimate = 1.13 [1.05, 1.22]; Fig. 2F; Fig. 4).

**Figure 2.**
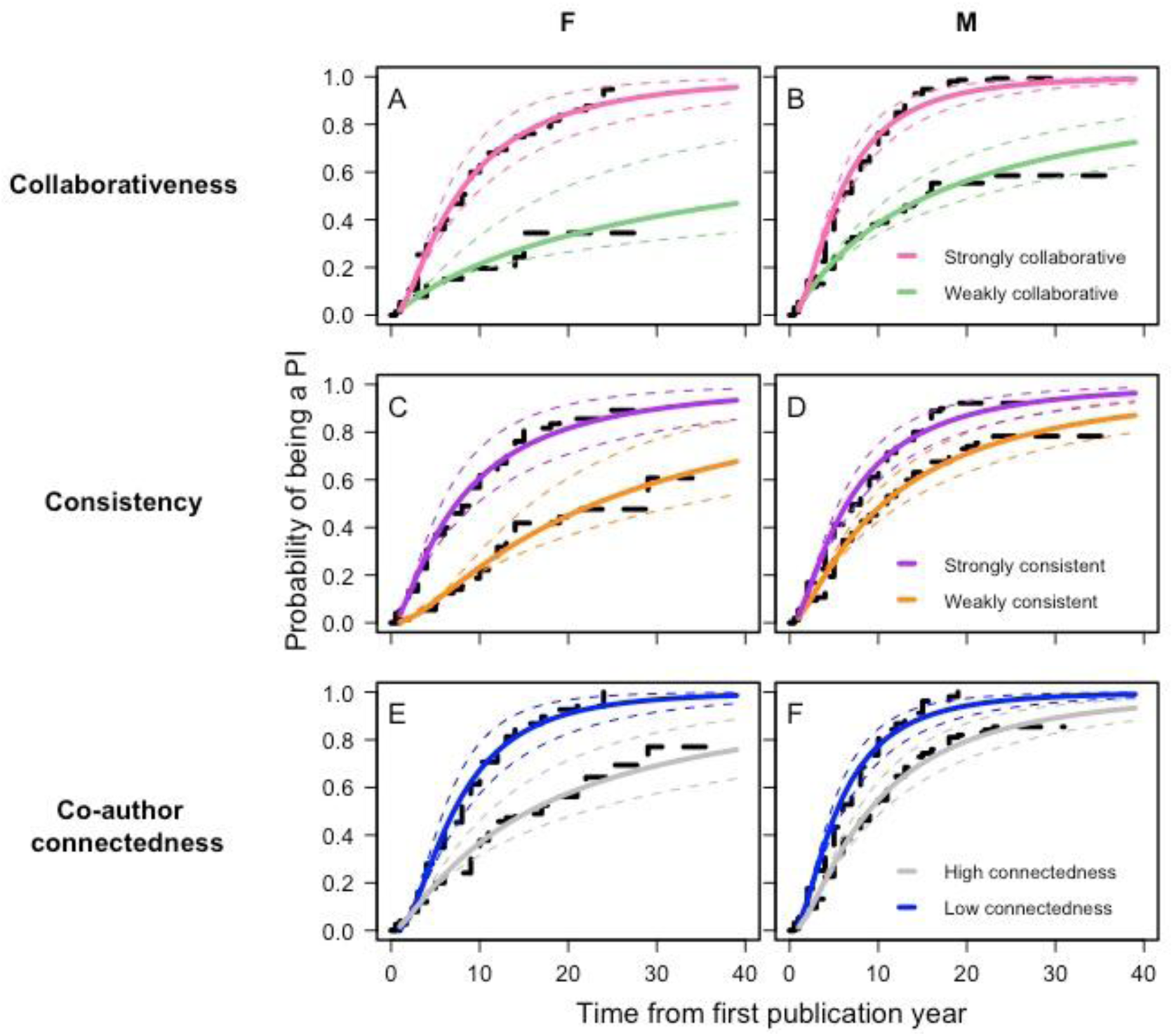
The impact of publication output-corrected egocentric network metrics (collaborativeness [n = 298 F; n = 637 M], consistency [n = 298 F; n = 637 M], co-author connectedness [n = 296 F; n = 634 M]) on the time taken to achieve PI status (expressed as difference in years between the first publication and the second last-author authored publication), for female and male focal authors. Analyses were conducted on continuous data, but the two lines visualised in each plot represent the 25^th^ and 75^th^ quartile of the data for each variable. Black dashed lines represent Kaplan-Meier survival functions, while the coloured lines represent model predictions (solid lines) with 95% confidence intervals (dotted lines).

### Career length

Collaborativeness positively predicted career length in both genders. Focal authors that published with a greater number of co-authors (given their number of papers) were more likely to continue publishing than focal authors with fewer co-authors than average (Fig. 3A, B). There was a significant interaction between this author sociality metric and gender, with the effect of collaborativeness on career length being nearly double for female focal authors compared to male focal authors (interaction term: AFT, estimate = 0.51 [0.39, 0.66]; gender-specific models: female focal authors, estimate = 3.93 [3.16, 4.89]; male focal authors, estimate = 2.33 [2.02, 2.68]; Fig. 4). Thus, the impact of collaborativeness on career length, as for time to PI, was greater for female focal authors than it was for their male counterparts.

**Figure 3.**
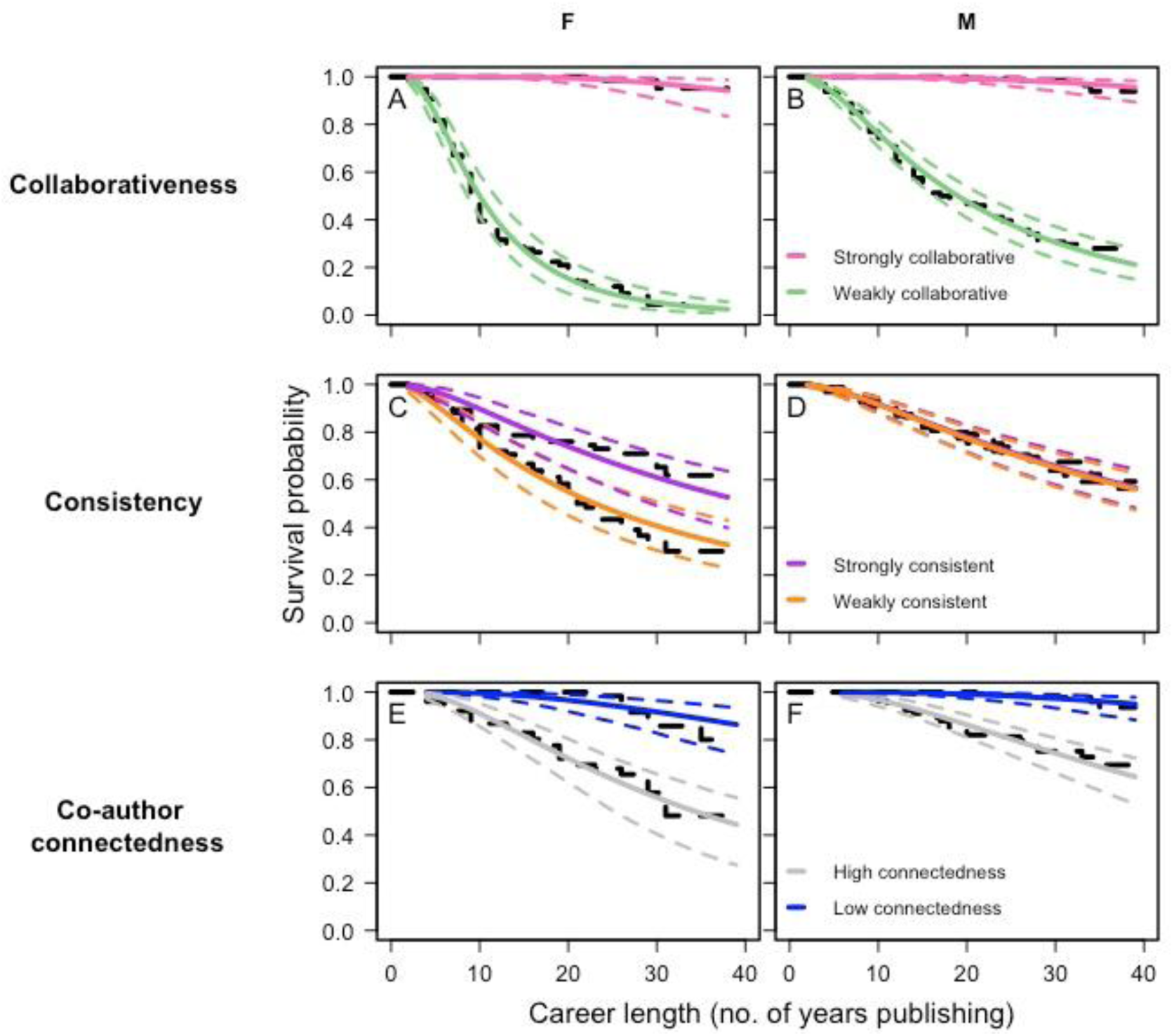
The impact of publication output-corrected egocentric network metrics (collaborativeness [n = 298 F; n = 637 M], consistency [n = 298 F; n = 637 M], co-author connectedness [n = 296 F; n = 634 M]) on career length (expressed as publication years), for female and male focal authors. Analyses were conducted on continuous data, but the two lines visualised in each plot represent the 25^th^ and 75^th^ quartile of the data for each variable. Black dashed lines represent Kaplan-Meier survival functions, while the coloured lines represent model predictions (solid lines) with 95% confidence intervals (dotted lines).

The effects of consistency (how frequently a focal author published with the same co-authors, given their number of papers) differed between female and male focal authors (interaction term: AFT, estimate = 0.77 [0.64, 0.93]; Fig. 4). For female focal authors there was a positive, but non-significant effect of consistency on career length, with increased consistency resulting in longer careers (AFT, estimate = 1.16 [0.99, 1.35]; Fig. 3C). For male focal authors, greater consistency negatively and significantly influenced career length (AFT, estimate = 0.89 [0.80, 0.99]; Fig. 3D). Male focal authors who published more consistently with the same co-authors were on average 11% less likely to still be active in science compared to those with lower consistency scores.

**Figure 4.**
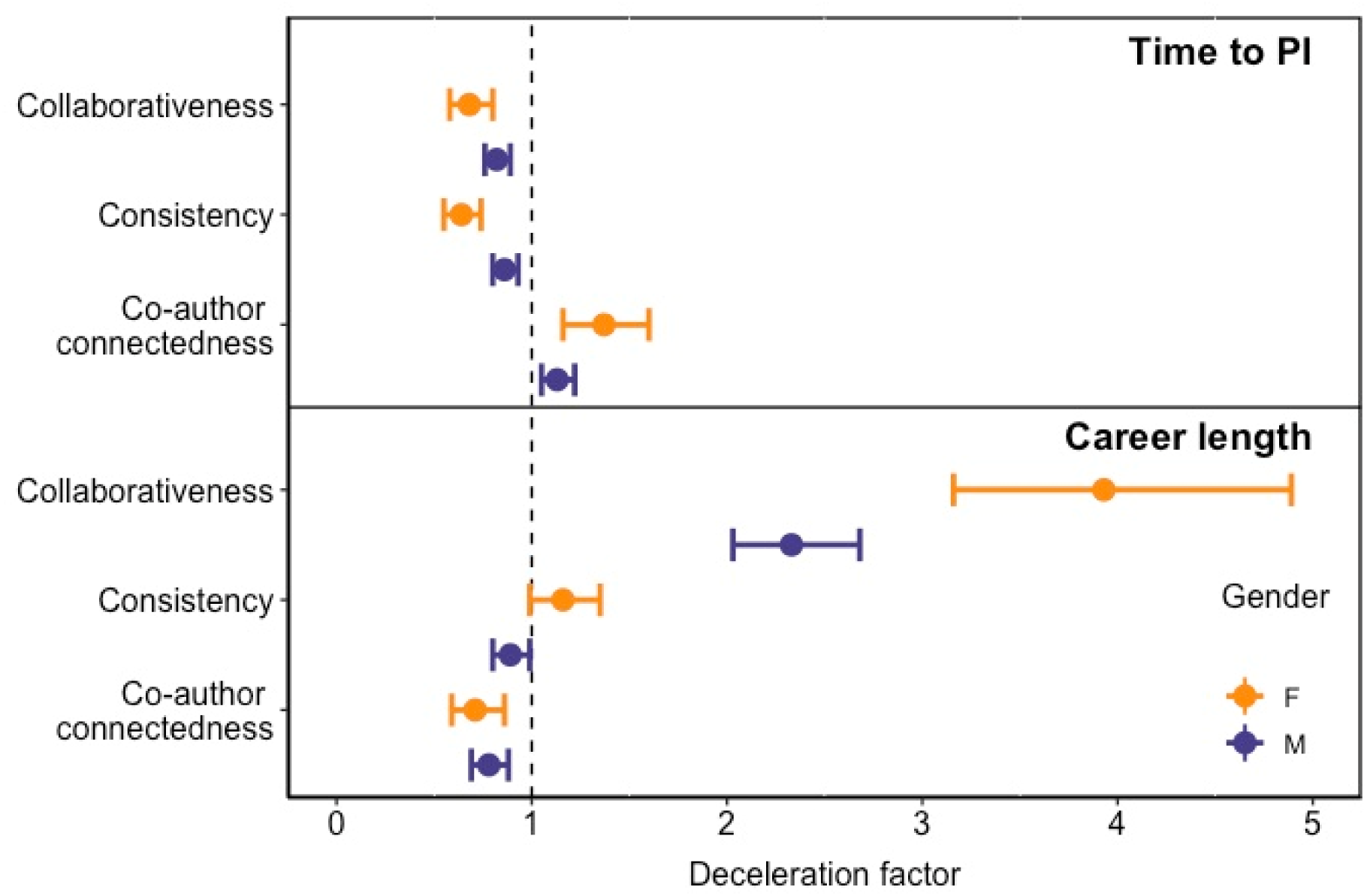
The relative effect size of publication output-corrected egocentric network metrics (collaborativeness [n = 298 F; n = 637 M], consistency [n = 298 F; n = 637 M], co-author connectedness [n = 296 F; n = 634 M]) on the time taken to achieve PI status, and career length, for female and male focal authors. The deceleration factor represents the AFT model estimate (based on separate models for female and male focal authors), and thus provides an indication of the magnitude and direction of effect of a particular egocentric network metric on our two measures of scientific performance. Filled symbols indicate the estimate, with error bars representing 95% confidence intervals. Estimates to the left of the dotted vertical line (<1.0) indicate a reduction in the time to the parameter of interest (i.e. reduced time to PI, or career length), and vice versa for estimates >1.0. Estimates with confidence intervals overlapping 1.0 are non-significant while, within a particular social metric, overlapping confidence intervals imply that the estimates do not differ significantly between females and males.

A higher co-author connectedness (having many co-authors that repeatedly appear together on a focal author’s papers) significantly decreased the chance to remain scientifically active, but this was not significantly different between the sexes (interaction term: AFT, estimate = 1.11 [0.88, 1.38]; co-author connectedness: estimate = 0.76 [0.69, 0.84]). An increase of one unit in co-author connectedness score decreased the chance of remaining in science by approximately 24% (Fig. 3E, F; Fig. 4).

With the exception of the strongly positive influence of collaborativeness on career length, all three measures of author sociality were broadly similar in the magnitude of their effect on time to PI or career length. All three measures of author sociality, however, had more pronounced effects for female focal authors than for their male counterparts (Fig. 4).

### Directionality of effect: Does publishing behaviour at early career stage predict future academic survival?

Network metrics extracted from focal authors based on their first ten years of publishing showed very similar patterns in their relationship with scientific career survival to those based on analysis of the entire dataset (Table S2). This was the case both for the direction and the relative magnitude of any effect, and is in support of the hypothesis that network metrics drive career length, rather than the other way round (see Methods: Does author sociality affect scientific career length, or vice versa?). Extent of sociality during a focal author’s early career therefore predicted their future survival. Importantly however, in contrast to our findings based on the complete dataset, results for collaborativeness and co-author connectedness were independent of gender. This implies that any differences in how these two measures of author sociality affect the survival of female and male focal authors only arise after their first ten years of publishing, which corresponds approximately to the median time that focal authors in our dataset become PIs.

## DISCUSSION

While the role of research collaborations in promoting publication output and citations has been previously reported ^48,65–69^, our study is the first to explicitly demonstrate the benefits of collaborative research to the progression and length of academic careers in science. Our findings reveal that scientists that collaborated more, had stronger connections to their co-authors, and that published with a wide diversity of relatively unconnected co-authors were more likely to become PIs, and became one more quickly. Being more collaborative and having less connected co-authors also increased the likelihood of remaining active in science for longer. Thus, the main finding of our study is that collaboration with a diverse community of co-authors is beneficial to career prospects and survival in science. The positive effects of collaborating early with eminent scientists have previously been reported ^70^ but our study suggests that such benefits arise from collaboration in general, and not only with leading scientists, or at early career stages.

Similar to studies from across STEM subjects that highlight gender differences in productivity and career advancement ^8,71,72,76,77,79,80,93–95^, we found that female researchers published fewer papers (equivalent to a 58.5% gender gap in productivity in our dataset), were less likely to become a PI, took longer to become one, and had shorter careers on average than male authors. Shorter career lengths and higher dropout rates explain a large proportion (but not all) of the productivity and impact differences that female researchers suffer compared to their male colleagues ^76^. Our study highlights how such negative effects for women could be magnified: if reduced career length and lower productivity also restrict opportunities for enhancing sociality by establishing new collaborations, then this would further contribute to less favourable career outcomes for female academic scientists.

Female focal authors were less collaborative than men (i.e. had fewer unique co-authors), as has been previously documented ^71,73,74,96^ (but see ^72^), and this result remained after controlling for productivity differences between female and male focal authors (see Methods: Accounting for the effect of publication number on social metrics). This is in contrast to the findings of ^71^, who found that the difference in co-author numbers between genders could be fully explained by the lower publication rate and shorter career lengths of women scientists. Nonetheless, collaborative style ^97–99^ could be equally as important as the number of collaborators for determining the value of collaborations to a researchers’ career. Furthermore, the publishing networks of female and male focal authors showed similar levels of consistency and connectedness, suggesting that any inherent differences in sociality between female and male researchers are likely small. Given this, a striking finding of our study was that the effects of author sociality on career progression and length were consistently stronger for female than for male focal authors. One explanation for this may be that female researchers are judged more on the basis of whom they work with than are male colleagues, making their collaborations all the more important ^22,100–102^. The gender composition of a researchers’ network may also be more important for female researchers than for male scientists. For example, more prestigious roles within collaborative projects, including senior and corresponding authorships, are often assigned to the detriment of women, potentially causing their contributions to research to be undervalued ^103,104^. Having a female senior author on a manuscript, however, increases both the overall proportion of women authors, and the probability of a female first author ^104^. Similarly, having an inner social circle composed predominantly of women predicts women’s leadership success ^105^. Alternatively, since researchers are more likely to publish with colleagues of the same gender than is expected by chance ^99,106^, gender preferences in co-authors may further reinforce gender differences in career progression and length. This effect could be further compounded since high-performing (and therefore ideal collaborator ^102,107^) male academics employ relatively fewer women, and thus provide fewer associated collaboration opportunities ^24^.

Consistency was the only measure of co-author sociality that had a negative effect on any of the performance measures we assessed, and this only applied to career length for male focal authors. Previous research indicates that the majority of collaborations between researchers are weak and transitory ^108^. However, those co-authors that a scientist collaborates with repeatedly over their career – and that would therefore tend to raise the consistency score of a focal author in our dataset – contribute significantly and positively to above-average productivity and increased citation rates ^108^. These are both factors that would likely promote career advancement and length. Indeed, we found that higher consistency positively influenced career progression in both genders. It is therefore unclear why being more consistent in who you publish with should reduce career span for men, particularly when there was no significant effect of consistency on career length for female focal authors. ‘Star’ collaborators – those eminent scientists that stand out because of their high funding rate, citation rate or prize-winning ability – can significantly enhance the productivity and citation rate of their co-authors ^102,107^. Closer investigation of exactly who it is that female and male focal authors collaborate with, and how often, may shed more light on this peculiar effect of consistency on career length.

In addition to providing insights into how sociality differentially affects the careers of female and male scientists, our study also highlights the value of applying concepts from ecological and evolutionary studies of sociality to understand survival in science. Indeed, we find it intriguing that, despite building social networks based solely on publication affiliations, our results show strong parallels with studies across a range of species and contexts that investigate the role of sociality in enhancing performance and longevity ^31,38,41,43^. These studies often emphasise the importance of early life conditions ^33,35,109,110^, and our findings point to a similar effect. Social network metrics calculated based on the first ten years of a focal authors’ career were predictive of their overall career length, suggesting that the approach to collaboration and publishing that a scientist takes early on in their career has important implications for their future success ^70,111^. Junior scientists are typically more likely to be pursuers of collaborations, while senior researchers are net attractors, having new collaborations proposed to them ^108^. Our findings suggest that for scientists early in their career, being an enthusiastic pursuer of collaborations is beneficial in the long term, especially since the tendency to collaborate increases with career length ^98^, and how collaborative a scientist is predicts their productivity better as their career progresses ^112^. With the exception of consistency, we found no effect of gender on how early career levels of sociality predicted career length, in contrast to the clear gender asymmetry apparent in the entire dataset. We suggest that the gender-specific effects of author sociality may arise only around the time that a researcher becomes a PI.

Although our study focused specifically on contributors to International Society for Behavioral Ecology conferences, the range of researchers that attend these conferences is wide (see Methods: Identification of focal authors), and we expect our findings to be applicable across ecology and evolutionary biology. Over the course of their careers, the researchers in our study published papers on a wide diversity of subjects in these disciplines, and beyond. Furthermore, both the positive effects of sociality that we observe, and the issues surrounding collaborative working, are unlikely to be subject-specific. Regardless of discipline, finding and establishing new collaborations is not always straightforward, and working more collaboratively may require overcoming barriers, such as having a small peer network, or lacking access to funding ^113^. This may be a particular challenge for female researchers, who represent less of the workforce as seniority increases ^9^, and especially since male researchers typically prefer to collaborate with other men ^99,106^. Researchers may also have concerns over the risk to reward ratio of forming new collaborations ^113^. A larger team of co-authors on a research project entails sharing any reward for the work amongst a larger group of colleagues. Our findings imply that this should not deter researchers from seeking out new collaborations. Rather, more collaborators may increase the number of new networks that a scientist and their work can be introduced to. Indeed, collaboration is frequently mentioned as an important factor in scientists’ own reflections on their success ^114^ in ^115^. In conclusion, our results suggest that all researchers – but particularly those that are female – can enhance their career progression and survival in science by collaborating widely and repeatedly. Creating research environments that encourage collaboration – across disciplines and institutes, among all career levels, and especially between the genders – will lead to greater and more rapid scientific advances ^20^, but can also assist in reducing the gender gap in science.

## METHODS

### Identification of focal authors

Our subjects were contributors to the International Society of Behavioral Ecology (ISBE) biennial meetings in 1992 (Princeton, USA, 766 participants), 1994 (Nottingham, UK, 521 participants) and 1996 (Canberra, Australia, 565 participants). ISBE conferences attract researchers working in the field of behavioural ecology, but also in related areas including animal behaviour, evolutionary biology, population biology, physiology, and molecular biology (www.behavecol.com), and contributors therefore represent a broad spectrum of research fields. We chose these three particular conferences because they occurred long enough ago that any participants who were at the start of their career in the early 1990s would by now have had sufficient time to reach PI status, should they have tried to do so (how we define this is specified below). We also considered it likely that, particularly for junior or early career researchers, for whom access to funding might be more limited, most delegates would only be attending an international conference if they had a talk or poster to present. Thus, contribution to a conference likely indicated active participation in science at that time. Conferences are a common place for scientists to meet and find existing and new collaborators ^116^, and in the 1990s the ISBE conferences attracted many of the most active researchers in behavioural ecology and related fields, and also many young students embarking on a career in this field (pers. comm. Prof. Lotta Sundström and Prof. Nick Davies, Feb. 2020). Finally, these conferences spanned three continents, thereby maximising the global coverage of our sampling. Given these reasons, we are confident that our initial study cohort captures as large and as representative a sample of the behavioural ecology community at that time as we could hope to achieve.

A total of 1469 unique names were listed in the conference booklets for all three conferences. There was no consistent format to how information on conference participation was presented, and so all names were included in our initial pool of subjects, regardless of whether they were listed as a delegate, a registrant, a participant, were presenting a talk or a poster, or were a co-author on a presented talk or poster. Whilst our starting list may include some researchers who did not physically attend any of the conferences (we have no way of knowing this), our aim was to include all those who may nonetheless have contributed to the scientific content of these conferences in some way, and thus could be identified as potentially being scientifically active. Our next steps in refining this list (see ‘Bibliometric procedures’ and ‘Restrictions’, below) further identified those who we considered to actually be active in science. Where possible, we recorded the gender of conference contributors as male or female, based on their first name and/or online profile, and noted the continent that they were associated with at the time of their first conference registration. We then imported this list of individuals and associated attributes into R version 3.5.0 ^117^ in which all further data processing, analyses and visualisations were carried out.

### Bibliometric procedures

We used the ‘rscopus’ package ^118^ to identify all those individuals in our starting pool of conference participants that also existed in the academic database Scopus (www.scopus.com), which claims to be “the largest abstract and citation database of peer-reviewed literature”. We filtered our list to only include those contributors that had a Scopus Author Identifier (SAI), indicating that they had published at least one scientific article. Subsequently, we manually searched Scopus by author name (first initial and surname, gender anonymised), further restricting our dataset of potential authors to those that had published at least one paper that could be considered as relevant to the field of behavioural ecology, as judged from the title of the paper, the contents of the abstract, the research specialisation(s) of the authors, and the journal of publication. We also combined the publication records of any identified authors who had multiple Scopus profiles, meaning that their output could be found through different SAIs. From our initial list of 1469 conference contributors, 108 did not have a SAI, suggesting that they had never published a peer-reviewed paper. In a further 125 cases with multiple SAIs, it was not possible to determine which was correct, either because an ambiguous author name yielded too many search hits, and/or because it was unclear from the publication history. This left us with a dataset of 1236 conference contributors who had all published at least once, and whom we will now refer to as focal authors.

We downloaded the author list, title, publication year and journal of all publications for all focal authors (as of September 2018) using the ‘bibliometrix’ package ^119^, in conjunction with Scopus API keys (www.api.elsevier.com) and manual searches (see supplementary methods S2 for more information). To minimise the chance of name synonymy – the same author appearing under slightly different names in our dataset – the first name of each focal author in our dataset was reduced to the first initial, and any accents were removed ^120^. In four cases, two focal authors had the same surname and initial, and were thus marked with a unique number to ensure they were recognised as separate individuals in the analyses.

### Restrictions

We applied several restrictions to our initial dataset of focal authors. We excluded focal authors that only published single-author papers (n = 9 authors), or had published fewer than three papers in total (n = 91 authors), as it was not possible to construct meaningful social networks for these focal authors. We also excluded focal authors with more than 400 papers (n = 6 authors), as they stood out as outliers (Fig. S1A). To produce a homogenous cohort, and to minimise issues arising from changes in publishing patterns over time (e.g. increasing numbers of co-authors on papers ^30,120–122^) we further restricted our dataset to those focal authors who published their first paper in 1980 or later (thereby excluding n = 192 authors; Fig. S1B). For those focal authors that remained in our dataset, we then refined their output so that we only considered articles that were published in academic journals. After manually correcting for errors or discrepancies in journal titles, the following types of records were all excluded (6% of total output, affecting 394 authors): books and book chapters, trade journals, conference papers and proceedings, editorials, errata, and short surveys. Finally, any articles with 20 or more authors (1% of all papers in the dataset; affecting 330 authors) were excluded from analyses, since these were extreme outliers, and not representative of the general publishing norms within our dataset (Fig. S1C). This left a final sample size of 938 focal authors with a total of 53,279 papers. We checked whether any of these exclusions disproportionately affected either gender, given the underlying proportion of female and male focal authors in our initial dataset. This was not the case (see supplementary methods S3).

### Egocentric network construction

Having defined our final dataset of focal authors we then generated an egocentric social network for each focal author, based on their individual list of publications and co-authors. Egocentric networks have the subject of interest (here, the focal author) as the central node (the ego) ^123^ and include connections to and among social contacts (‘alters’, or co-authors in this case) that are adjusted by the strength of the association between ego and alter (here equivalent to the number of co-authored papers that focal author and individual co-authors share; Fig. S2) ^124^. We calculated social network metrics by transforming the publication record of each focal author into a binary ‘group by ID’ data structure with the ‘asnipe’ package ^125^, in which each row represented a paper, and every column was an author. This was then converted into a weighted adjacency matrix in the ‘asnipe’ package, from which we created weighted graphs with the ‘igraph’, ‘intergraph’ and ‘network’ packages ^126–128^.

### Egocentric network metrics

We measured three metrics from each focal author’s egocentric network. First, we measured network size, or ‘degree’, which corresponds to the total number of co-authors in the focal author’s network. A higher degree indicates that an individual is associated with a greater number of unique co-authors. Next, we extracted ‘mean tie strength’ (tie strength hereafter), measured as the total number of times a focal author has published with her/his co-authors across all their papers, divided by the number of unique co-authors. Tie strength in this context thus provides an average measure of how many times a focal author publishes with the same co-authors. Finally, we measured the ‘global clustering coefficient’ of each focal author’s egocentric network, using the transitivity function in the ‘igraph’ package ^126^. This measure, ranging from zero to one, describes how connected, on average, all authors in a focal author’s network are, considering only the focal author’s papers ^129^. A high connectedness – signified by a clustering coefficient close to one – indicates that the same co-authors appear together on many of the focal author’s publications. This might be the case, for example, for a professor and her immediate group of postdocs and PhD students. It is important to note that the clustering coefficient implies nothing about how many papers co-authors may have published together without the focal author. We could not calculate a clustering coefficient for five focal authors in our dataset who had a maximum of one other co-author per publication.

### Accounting for the effect of publication number on social metrics

Unsurprisingly, all three metrics were significantly influenced by how many publications a focal author produced (degree: Generalised linear model (GLM), Poisson, estimate ± SE = 0.009 ± <0.001, *z*_936_ = 233.4, *p* < 0.001; tie strength: GLM, Gaussian, log10-transformed; estimate ± SE = 0.002 ± < 0.001, *t*_936_ = 13.85, *p* < 0.001; global clustering coefficient: GLM, Binomial; estimate ± SE = -0.006 ± <0.001, *z*_931_ = -595.5, *p* < 0.001; Fig. S3). That is, focal authors with more papers tended to have more co-authors overall (a higher degree), to publish more often with the same co-authors (a greater tie strength), and to have co-authors that published less often with each other (a lower global clustering coefficient). These effects were further compounded by clear gender differences in our dataset in terms of the number of papers that female and male focal authors published. Overall, female focal authors published significantly fewer papers than male focal authors (GLM, Poisson, estimate ± SE = 0.54 ± 0.01, *z*_933_ = 50.81, *p* < 0.001; F: 38.46 ± 2.30 papers, median = 23.5 papers, range: 3 – 234 papers; M: 65.71 ± 2.42 papers, median = 51 papers, range: 3 – 375 papers), equivalent to a gender gap in productivity of 58.5%. As a consequence, female focal authors had significantly fewer co-authors (i.e. smaller networks and thus lower degree values; GLM, log10-transformed, estimate ± SE = 0.57 ± 0.08, *t*_933_ = 6.88, *p* < 0.001; Fig. S6A), published with their co-authors less frequently (GLM, log10- transformed, lower tie strength; estimate ± SE= 0.08 ± 0.02, *t*_933_ = 3.69, *p* < 0.001; Fig. S6B), and published with co-authors that were less connected (global clustering coefficient; GLM, binomial, estimate ± SE = -0.30 ± 0.002, z_928_ = -170.7, *p* < 0.001; Fig. S6C), compared to those of male focal authors. To account for these strong and gender-biased effects of publication number, for each focal author we therefore regressed all three metrics against their number of publications and used the residuals of these three models (hereafter, degree residuals, strength residuals, and clustering residuals, respectively) for all further analyses. All residuals were standardised by subtracting the mean and dividing by the standard deviation.

We used degree residuals to define a focal author’s ‘collaborativeness’, which can be considered a measure of how many unique co-authors a focal author publishes with, corrected for the number of papers they have published. Collaborativeness was analysed as a continuous variable. Thus, for a given number of published papers, a focal author with a positive collaborativeness score can be considered to be more collaborative than the average (i.e. publishes with a higher number of co-authors), while a focal author with a negative collaborativeness score is less collaborative than the average.

We used degree residuals again, together with strength residuals, to obtain a measure of how frequently a focal author published with the same co-author(s). We term this ‘consistency’. We extracted the residuals of a linear regression model with non-constant variances (‘lmvar’ package ^130^) that tested the effect of the degree residuals on the strength residuals (estimate ± SE = -0.36 ± 0.02, z_4_ = -15.60, *p* < 0.001). Consistency was treated as a continuous variable. Focal authors with a positive consistency score published more often with the same co-author(s) than the average for focal authors with the same number of papers, and vice versa for those with a negative consistency score.

Finally, we used the clustering residuals to define the co-author connectedness of a focal author. Focal authors with a positive co-author connectedness score publish with co-authors that are more highly connected than the average for other focal authors with the same number of papers. By contrast, a negative co-author connectedness score would indicate that, compared to the average for focal authors with the same number of publications, a focal author publishes with co-authors that have rarely or never published with each other.

Thus, for each focal author, we ended up with three new metrics, adjusted for the number of papers published, that describe how social they were in their approach to publishing: their collaborativeness, their consistency, and their co-author connectedness.

### Measures of academic performance and accounting for authorship name changes

We looked at academic performance in two ways: i) Following the definitions in ^94^, we classified any focal author with at least three last-author publications as being a PI, and measured time taken to reach PI status (time to PI) as the difference in years between their first publication and their second last-author publication; ii) Scientific career length was calculated as the time from first until last publication in our dataset ^1^. For the purposes of this study, we therefore consider a scientific career to be one that focuses on research and publication, and that can be quantified solely according to the metric of publication output. We fully recognise that there are other equally valid and important careers and means of contributing to science that are not research-focused and that cannot be quantified in this manner ^131^. Our definition of career length relies on correctly identifying the publications of focal authors. This task becomes more complicated if an author changes the name under which they publish during their career, as might be the case particularly for female focal authors (e.g. due to marriage). Unfortunately, there is little published work on how name changes affect indexing and citation accuracy (but see ^132^ for a notable exception), or how indexing services cope with this issue. We checked whether author name changes could have resulted in erroneous categorisation of a female focal author in our dataset as having left science (i.e. stopped publishing; n = 126). To do this we manually searched the Internet for any webpages that might contain publication lists for these female focal authors, including Google Scholar and ResearchGate profiles, as well as professional and personal webpages. We looked for any indication of a change in publishing name, as well as checking the number of publications that a focal author had produced, and the date of their last publication. We identified only two female focal authors who had changed their publication name and thus were incorrectly assigned in our dataset as having left science. The records for these focal authors were updated accordingly.

### Statistical analyses

#### Data independence

Whole social networks suffer from non-independence because metrics calculated for one individual in a network are dependent on other members of the network ^133,134^. If unaccounted for, this may cause overestimation of true sampling variance, and overconfidence in results ^133^. Egocentric networks can also suffer from this problem if many individuals occur across multiple networks. We tested the independence of our egocentric networks by determining the proportion of networks in which focal authors also appeared as co-authors, and the proportion of papers that consequently occurred in multiple networks. Focal authors occurred on average in < 1% of networks as co-authors, and in a maximum of 5.5% (for one focal author only) of other networks (Fig. S4A). Papers occurred, on average, in less than 1% of all networks (Fig. S4B). Thus, the egocentric networks in this study are almost entirely independent of one another. Given this extremely low level of overlap, we have followed the approach of ^134^, and used standard statistical procedures in our analyses.

#### Effect of Gender

We used general and generalised linear models to investigate how gender affected the social metrics of interest after correcting for the number of papers published by each focal author. We looked at how gender explained time to PI and career length by constructing accelerated failure time (AFT) models, which are parametric survival models that provide a robust statistical approach to analysing survival data. They are based upon the survival curve, rather than the hazard function, as is the case for Cox proportional hazard models, another commonly used approach for statistical modelling of survivorship data ^135^. AFT models provide an intuitive summary measure that describes the extent to which the survival curve is shifted forward or backward by the effect of the variable of interest. The extent of this forward or backward shift is determined by the parameter *c*, also known as the deceleration factor, in the relationship *S*_1_(*ct*) = *S*_0_(*t*), where *S*_1_(*t*) is the survivorship of a cohort receiving a particular ‘treatment’ at time *t*, while *S*_0_(*t*) is the survivorship of a control cohort also at time *t*. If the ‘treatment’ (e.g. being more collaborative) increases survival, this has the effect of shifting the survival curve forwards, as represented by a value of *c* > 1 ^136^. A confidence interval around *c* can be calculated and, assuming that similar models are built, the relative effect of different treatments on survival can be compared. For an AFT approach to be valid, survival times are assumed to follow a parametric distribution. All AFT models in this study were run with a lognormal distribution, following visual examination of model fit, and comparison of log-likelihood and AIC values for models with different distributions. Residuals of AFT models were also examined for goodness of fit with the chosen distribution, and in all cases were found to be a good fit. Validity of assumptions for normality and homoscedascity for linear models were checked by visual inspection of residuals and normal probability plots. Results of linear models are presented as mean ± SE, unless stated otherwise, while we provide estimates of the deceleration factor, with 95% confidence intervals, for the output of AFT models, and with ‘female’ always set as the reference gender. Estimates from AFT models with confidence intervals overlapping 1.0 are non-significant. We ran AFT models using the functions survreg and flexsurvreg in the ‘survival’ and ‘flexsurv’ packages respectively ^137,138^. We used the ‘lme4’ package ^139^ to run general and generalised linear models.

#### Effect of author sociality on time to PI, and career length

We ran AFT models to evaluate whether collaborativeness, consistency and co-author connectedness predicted time to PI and career duration. We included gender, and the interaction between gender and the social metric of interest, in all models. Where there was a significant interaction between gender and a measure of author sociality, we re-ran models for each gender separately. When this interaction was not significant, models were re-run without the interaction term to obtain an estimate for the relevant measure of author sociality. We also calculated restricted mean survival times using the ‘survRM2’ package ^140^. A restricted mean survival time is a measure of average survival up to a specific time point. It is calculated from a survival curve by determining the area under the curve (AUC) and then comparing the AUCs for different survival curves (for example, to examine gender differences in survival up to a certain time ^141^).

In a first set of survival models we calculated the fraction of focal authors that became a PI within *x* years since their first publication year. Seven hundred and eight focal authors became PI (i.e. published at least three last-author papers) before the end of our sampling period (2018), leaving 230 focal authors (25%) that did not, and were therefore censored in this analysis. Eighteen authors produced their second last-author paper in the same year as their first paper, and because survival models would otherwise ignore these zero values, these were given dummy values of 0.5 years. Re-running survival analyses with these authors excluded from the dataset did not alter the outcome of our models. We ran binomial logistic regression models to ascertain the role of author sociality in the likelihood of focal authors becoming PIs.

In a second set of survival models we investigated the effects of the network metrics on focal authors’ career length (number of years publishing). The survival curves were calculated as the fraction of focal authors still publishing after *x* years since their first publication year. We assumed focal authors to have stopped publishing, and thus, from the perspective of this study, for their scientific career to have ended, if at least two years had elapsed since their last publication. This cut-off was chosen as the majority (80%) of publication gaps – years in which a focal author published no papers – were shorter than two years (Fig. S5). Note that posthumous publishing (i.e. publications that include the name of an author who is no longer alive) is not an issue for our analyses as such individuals still continued to contribute to science. Focal authors that were publishing in 2016 or later (68% of authors) were assumed to still be active, and were censored in the analyses. Female focal authors might have longer or more frequent career gaps if, on average, they are more likely to take time away from their careers than do male focal authors, for example due to maternity leave or extended child care duties. To account for this possibility, we repeated our analyses while taking a more conservative censoring approach that likely overestimated the number of censored subjects (still publishing in 2012 and 2014, respectively, and thus allowing for a gap between publications of up to 6 years). All analyses yielded highly comparable results (Table S1). We also determined whether the severity of our censoring approach (still publishing in 2012, 2014 or 2016) disproportionately affected our measure of the number of female or male focal authors that were considered to have left science. It did not (Supplementary methods S3).

#### Does author sociality affect scientific career length, or vice versa?

A positive correlation between any our three measures of author sociality and career length might arise if focal authors that have longer careers, or are in more senior positions in science, are deemed as more attractive to collaborate with, or have simply had more time to establish successful and enduring collaborations. To test the directionality of any relationship between our measures of author sociality and career survival we investigated whether a focal authors’ publishing behaviour in their early career predicted their future academic survival. If similar patterns arise between network metrics calculated at the early career stage, before focal authors have had time to establish their prominence and future academic survival, then we argue that this provides strong evidence that it is the network metrics measured here that drive career length, and not the other way round. To investigate this, we conducted a separate analysis of focal authors that started publishing the year prior to their first attendance at an ISBE conference or later (to produce a homogeneous cohort), and calculated the three author sociality metrics for up to their first ten publication years (*n* = 224 authors; 33.9% women; 3675 publications). We then ran AFT regression models to test the effects of network metric, gender, and their interaction, on career length, as previously described.

### Data and code availability

The data that support the findings of this study, in which focal author names are anonymised, will be available via Open Science Framework. Scripts for computational analysis of this data will also be available via Open Science Framework. We will also provide the code used to access publications available on Scopus for any given Scopus Author Identifier (SAI), as well as code for data wrangling, calculation of social network metrics and academic performance measures. For purposes of confidentiality, the names and SAIs of focal authors used in this study will not be made available. For reviewing purposes, data and code are available here: tiny.cc/vanderwal.

## ACKNOWLEDGEMENTS

We thank Neeltje Boogert, Alecia Carter, and Hannah Rowland for initial discussions about the project, Damien Farine for advice on early analyses, Wendy King for access to the ISBE archives, Rahia Mashoodh for assistance with dredging the Scopus API, Lauren Brent for advice with combining network and survival analyses, Chris Duncan for suggesting the use of AFT models, and Nick Davies, Lotta Sundström, and the Evolution, Sociality and Behaviour group at the University of Helsinki for valuable discussions about publishing, the history of behavioural ecology, and our results. J.E.M.vdW. was supported by a grant from the Association for the Study of Animal Behaviour (ASAB) awarded to R.T. R.T. was supported by an Independent Research Fellowship from the Natural Environment Research Council UK (grant number NE/K00929X/1) and a start-up grant from the Helsinki Institute of Life Science (HiLIFE), University of Helsinki. N.P.C.H. was supported by a Leverhulme Early Career Fellowship from the Leverhulme Trust.

## AUTHOR CONTRIBUTIONS

N.P.C.H. and R.T. conceived the study and all authors developed the concept. All authors collected the data. J.E.M.vdW. prepared the data for analysis and ran the social network analyses. N.P.C.H. carried out the AFT survival analyses. N.P.C.H. and J.E.M.vdW. prepared the initial draft and all authors edited and approved the manuscript.

## SUPPLEMENTARY MATERIALS

### Supplementary methods S1

Our final dataset included 298 women (32%) and 637 men (68%), and three focal authors for whom gender was unknown. Focal authors came from 39 different countries, with the majority from Europe (47%) or North America (35%). The mean ± SE number of unique co-authors per focal author was 81.09 ± 2.72 (median = 58; range = 1 – 628; Fig. S3), with each focal author producing, on average, 57.03 ± 1.85 papers (median = 42; range = 3 - 375 papers; Fig. S3). This equates to 3.86 ± 0.04 co-authors per paper (median = 3.72 co-authors; range = 1.11 – 11.11). Focal authors published with the same co-author(s) an average of 2.02 ± 0.02 times (median = 1.90; range = 1 - 7; Fig. S3). It took, on average, 9.34 ± 0.22 years for a focal author to become a PI, ranging from less than one year to 37 years (median = 8 years). The mean career lifespan was 24.67 ± 0.29 years (median = 26 years; range = 2 – 39).

### Supplementary methods S2

Due to the Scopus database and the Scopus Database ‘API’ Interface not being completely synchronised, we had to manually download publication details for 292 authors (24% of total) directly from the Scopus website, using the ‘search by Author ID’ function. To be sure that these two methods yielded identical results that could be merged, we extracted publication details using both methods for a random selection of 120 Scopus Author Identifiers. We found only two authors (< 2%) that differed in the number of papers assigned to them by the two search methods (one additional, and one fewer paper, respectively). We consider this validates combining data extracted via these two different methods.

### Supplementary methods S3

We applied a series of restrictions to our dataset of focal authors (see Methods for details), and for some survival models, we censored data. We checked to ensure that these restrictions and censoring decisions did not result in a gender bias through one gender being disproportionately excluded from the final dataset or particular analyses. The final dataset consisted of 32% women (n = 298). For every restriction in place (see below for a breakdown of numbers), we therefore confirmed that the percentage of women excluded or affected was not greater than 32%.

### Checking for gender bias in restricted data

- Focal authors that only published single-author papers: n = 9 authors excluded; 78 % women (7 women and 2 men)*
- Focal authors that published fewer than three papers in total: n = 91 authors excluded; 55 % women (50 women, 40 men, 1 unknown gender)*
- Focal authors with more than 400 papers: n = 6 authors excluded; 17% women (one woman and five men)
- Focal authors that published their first paper before 1980: n = 192 authors excluded; 11% women (22 women and 170 men)
- Non-journal articles: 0 authors excluded but 394 authors affected; 22% women (85 women and 309 men). For the women affected, 5.01% ± 0.66% (mean ± SE) of their total paper output were excluded, while for men this was 3.86 ± 0.20% of their papers (mean ± SE).
- Articles with more than 20 or more authors: 0 authors excluded but 330 authors affected.

* Although these restrictions are biased towards exclusion of female focal authors, they are both methodologically necessary in order to enable the construction of meaningful egocentric networks.

### Checking for gender bias in censored data for survival analyses

We applied three censoring cut-offs to our data, when determining the survival times of focal authors in our dataset: still publishing in 2012, in 2014 or in 2016. Using a more conservative censoring date did not change the proportion of female focal authors that were considered to have left science.

- 2012: 223 authors censored, 42.2% women (94 women, 127 men, 2 unknown gender)
- 2014: 252 authors censored, 42.1% women (106 women, 144 men, 2 unknown gender)
- 2016: 299 authors censored; 42.5% women (127 women, 169 men, 3 unknown gender)

**Figure S1.**
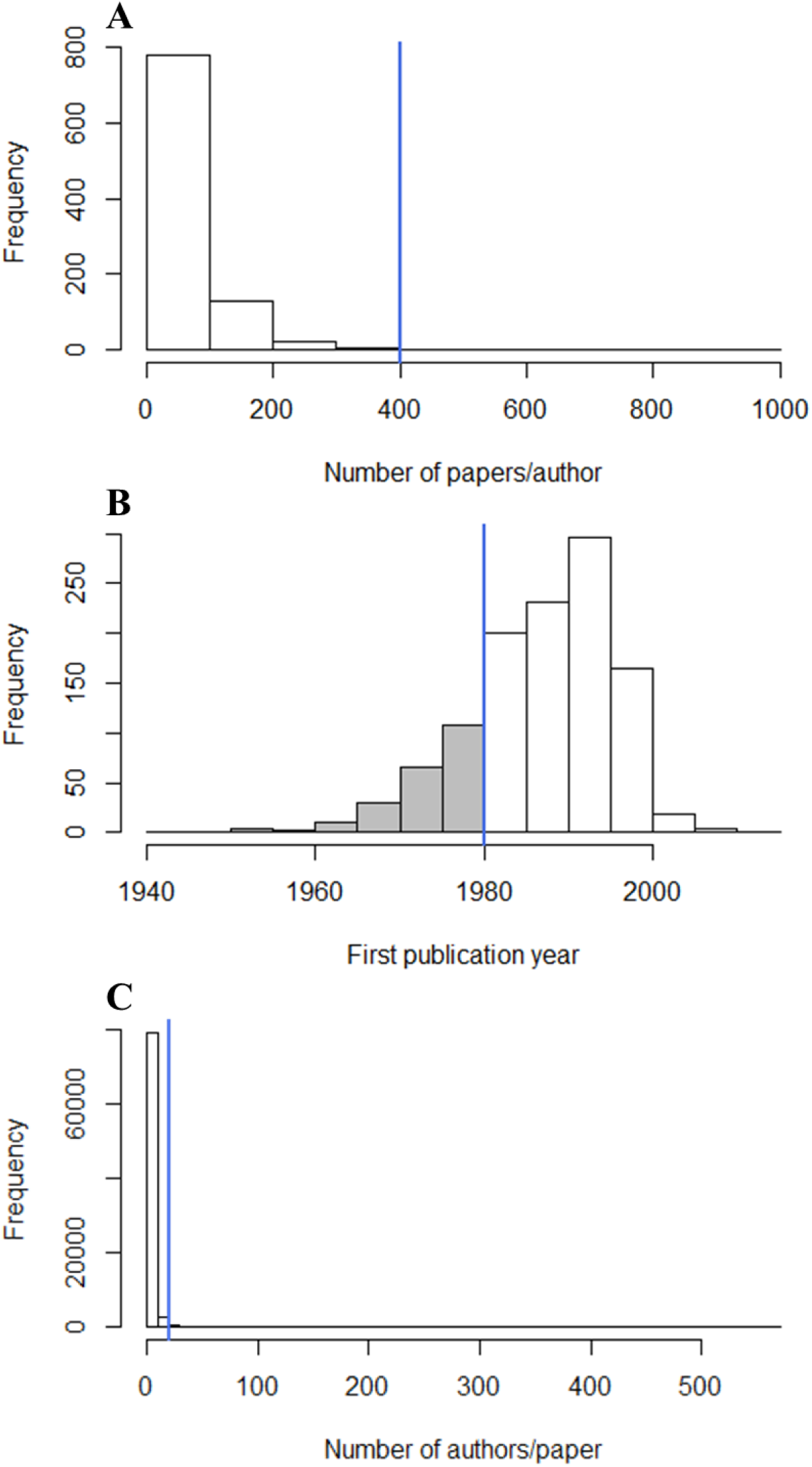
Visualisation of the restrictions that were applied to the final dataset. Blue lines indicate the implemented cut-off points and shaded areas indicate excluded data. (A) Papers with ≤ 400 authors were excluded. (B) Authors that started publishing on or before 1980 were excluded. (C) Papers with ≤ 20 co-authors were excluded.

**Figure S2.**
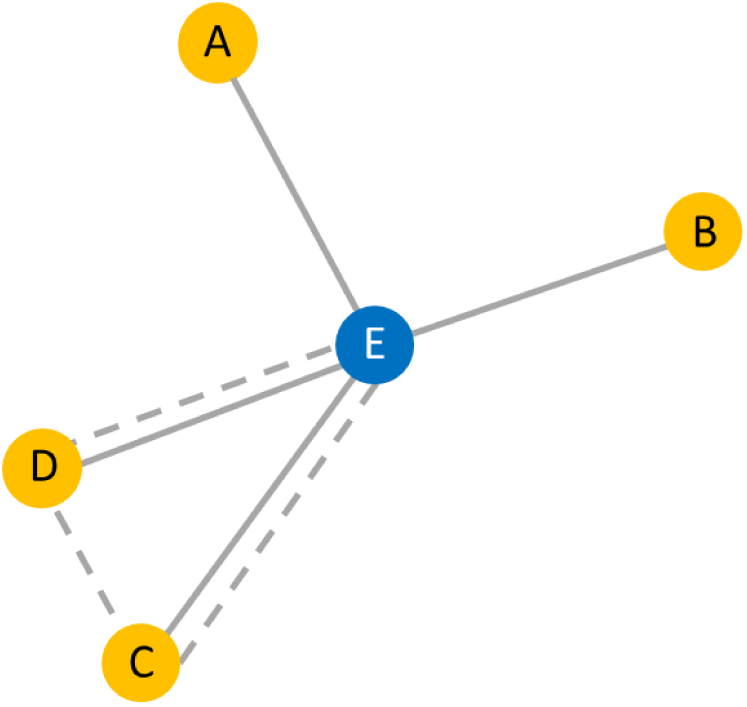
An anonymised egocentric network from our dataset with connections between the ego (focal author; blue dot, E) and co-authors (yellow dots, A-D) represented by grey lines. In this example, the focal author has published 5 papers in total: a single paper with each co-author, and a joint paper with co-authors C and D (represented by the dotted line joining E, C and D). Thus, focal author E has a network size or ‘degree’ of 5 (5 unique co-authors), and a tie strength of 1.25 (E has published, on average, 5/4 = 1.25 times per co-author). The global clustering coefficient for focal author E = 3 x number of triangles (one triangle: E-D-C)/number of all triplets (closed; E-D-C; D-E-C; D-C-E and open: A-E-B; C-E-B; D-E-B; C-E-A; D-E-A) = (3 x 1) / 8 = 0.375. After correcting for publication output (see ‘Accounting for the effect of publication number on social metrics’ in the Methods section) focal author E has a collaborativeness score of -1.27 (range: -6.62, 4.13), a consistency score of -1.28 (range: -2.61, 5.32), and a connectedness score of -0.08 (range - 5.42, 4.43). These are all negative values, implying that, compared to the average for other focal authors with the same number of publications, E is less collaborative, less consistent in who they publish with, and has co-authors that are relatively unconnected. Such a combination of social metric scores would suggest that focal author E probably did not become a PI, or else took a relatively long time to progress to this career stage, and may not have survived for particularly long in their career.

**Figure S3.**
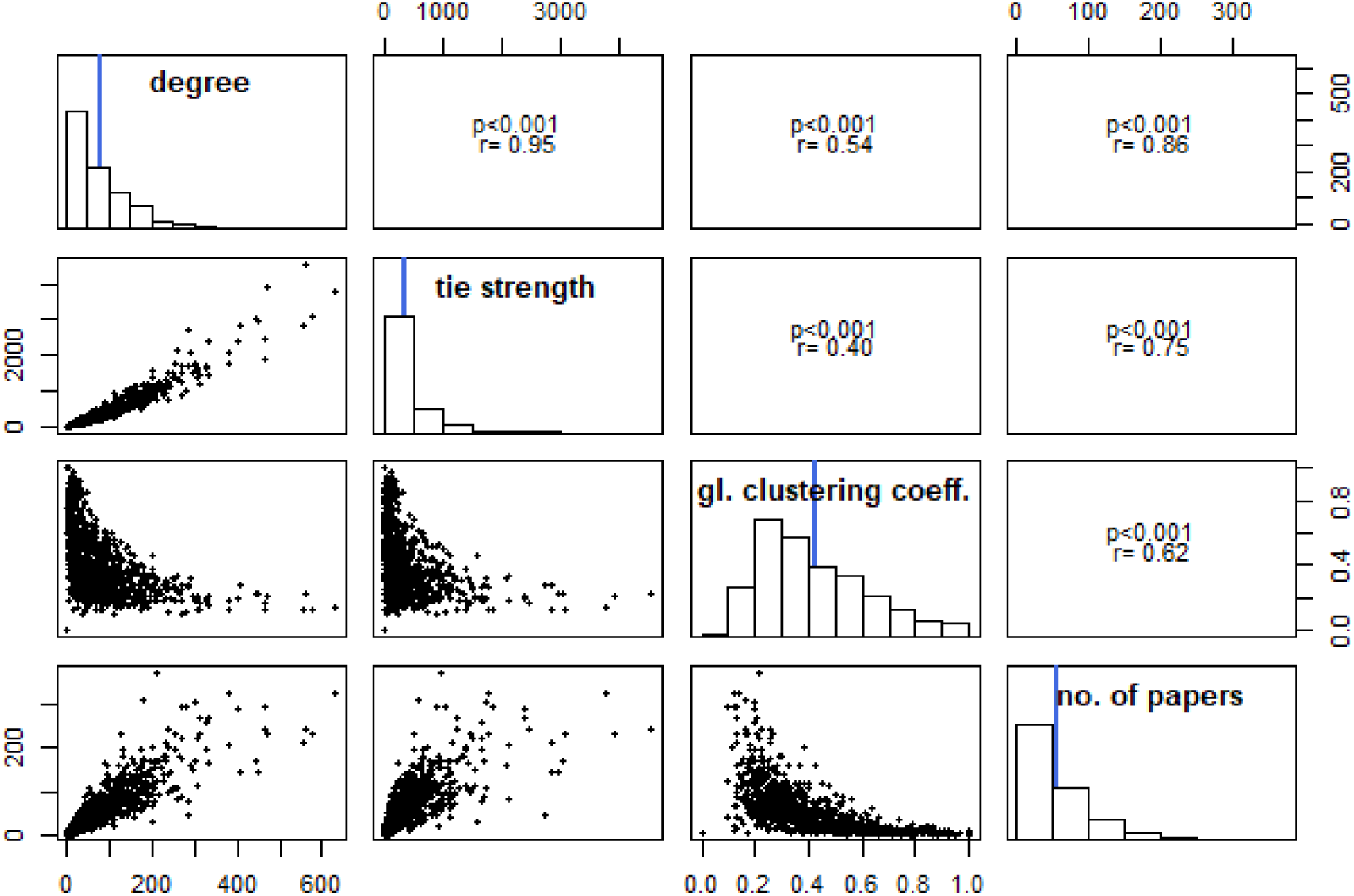
Correlation matrix and frequency histograms of the social network metrics measured, and number of papers (n = 938 focal authors). Blue lines in graphs indicate the mean value for each variable.

**Figure S4.**
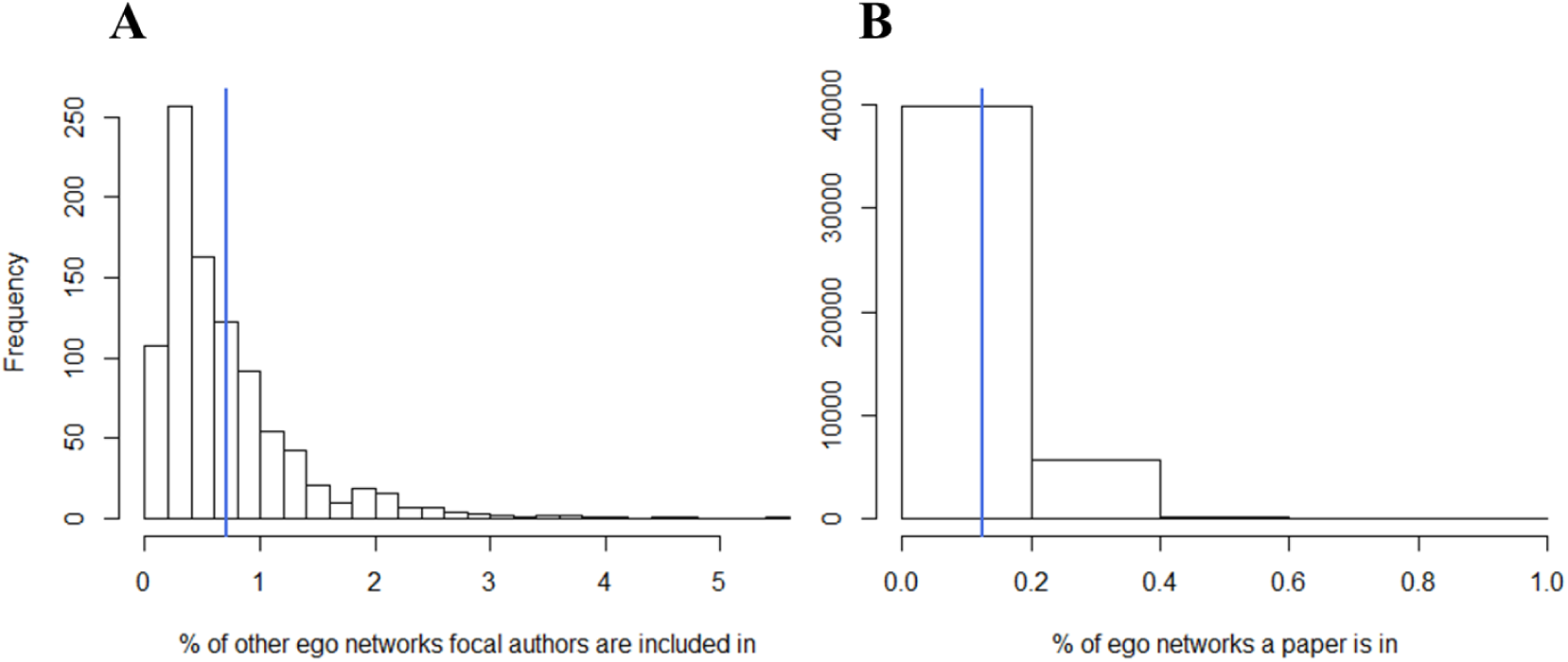
(A) A frequency histogram of the percentage of other ego networks that focal authors are included in. (B) A frequency histogram of the percentage of ego networks that individual papers appear in. In both panels the blue line represents the mean.

**Figure S5.**
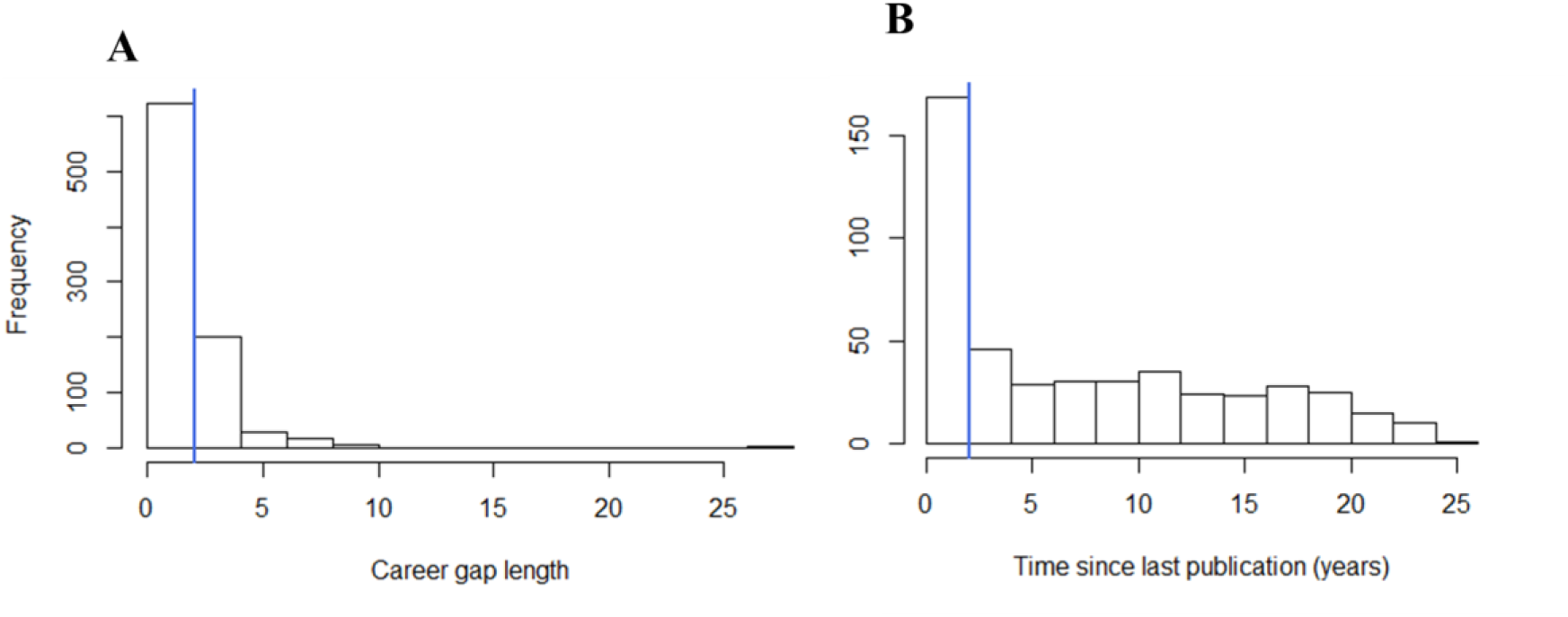
Censoring in the survival models. We censored focal authors (n = 938) that were still publishing in ≥ 2016, given that two years is the most common length of a career gap in the dataset, and the clear drop off in number of authors that published more than 2 years ago. We assume that all authors that published for the last time before 2016 were no longer actively contributing to science, and from the perspective of our survival analyses are thus ‘dead’. (A) Length of all career gaps (i.e. years between publications); (B) Years since last publication. In both panels the blue line indicates the two-year cut-off.

**Figure S6.**
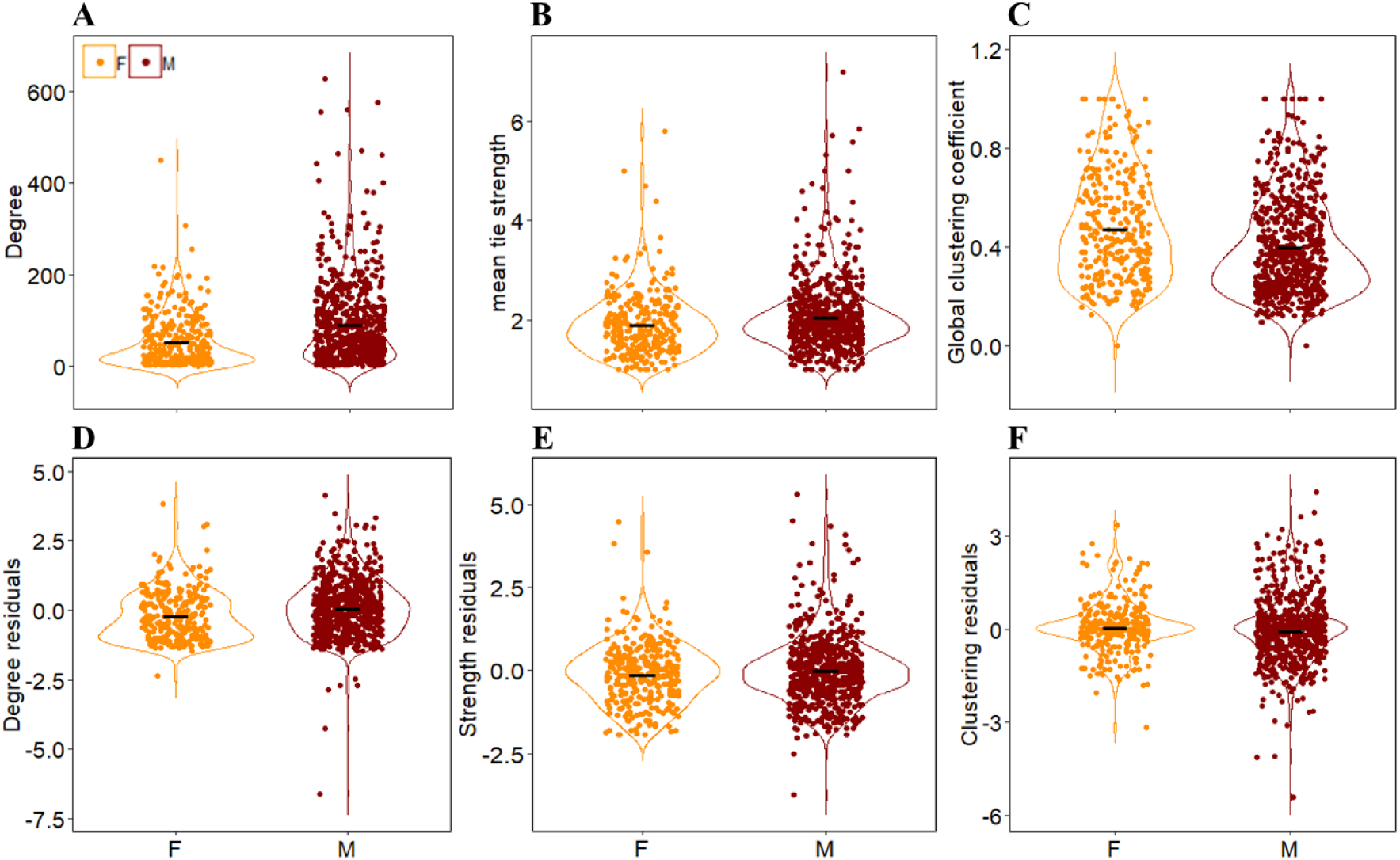
Significant gender differences in our social network metrics exist before correcting for number of papers (A-C). After correcting for publication output, males (n = 637) were found to be significantly more collaborative than females (n = 298) (D), while gender did not significantly affect consistency and co-author connectedness scores (E-F). Violin plots visualise the distribution of the data and its probability density. All raw (scaled) data points are displayed, and the black line indicates the mean.

**Table S1.** Output from AFT models testing the relationship between career length and each of the three measures of co-authorship sociality, in which we assumed authors to have stopped publishing when their last publication was before 2012, before 2014 or before 2016. Shaded estimates indicate non-significant relationships (95% confidence intervals overlapping 1.0).

|  |  | Estimate [95% CI] |  |  |
| --- | --- | --- | --- | --- |
|  | Term | 2012 | 2014 | 2016 |
| **Collaborativeness** | Interaction | 0.28 [0.18, 0.43] | 0.37 [0.26, 0.52] | 0.50 [0.38, 0.64] |
|  | Female | 7.70 [5.25, 11.29] | 5.83 [4.30, 7.92] | 4.00 [3.23, 4.96] |
|  | Male | 2.91 [2.38, 3.56] | 2.71 [2.27, 3.23] | 2.33 [2.02, 2.68] |
| **Consistency** | Interaction | 0.70 [0.55, 0.90] | 0.76 [0.61, 0.95] | 0.78 [0.65, 0.95] |
|  | Female | 1.14 [0.93, 1.40] | 1.12 [0.37, 1.35] | 1.13 [0.97, 1.32] |
|  | Male | 0.86 [0.76, 0.97] | 0.86 [0.76, 0.97] | 0.89 [0.80, 0.98] |
| **Co-author**<br>**connectedness** | Interaction | 1.06 [0.78, 1.43] | 1.03 [0.78, 1.35] | 1.13 [0.90, 1.41] |
|  | Female | 0.72 [0.56, 0.95] | 0.74 [0.59, 0.94] | 0.70 [0.58, 0.85] |
|  | Male | 0.77 [0.65, 0.90] | 0.76 [0.66, 0.88] | 0.78 [0.69, 0.88] |

**Table S2.**
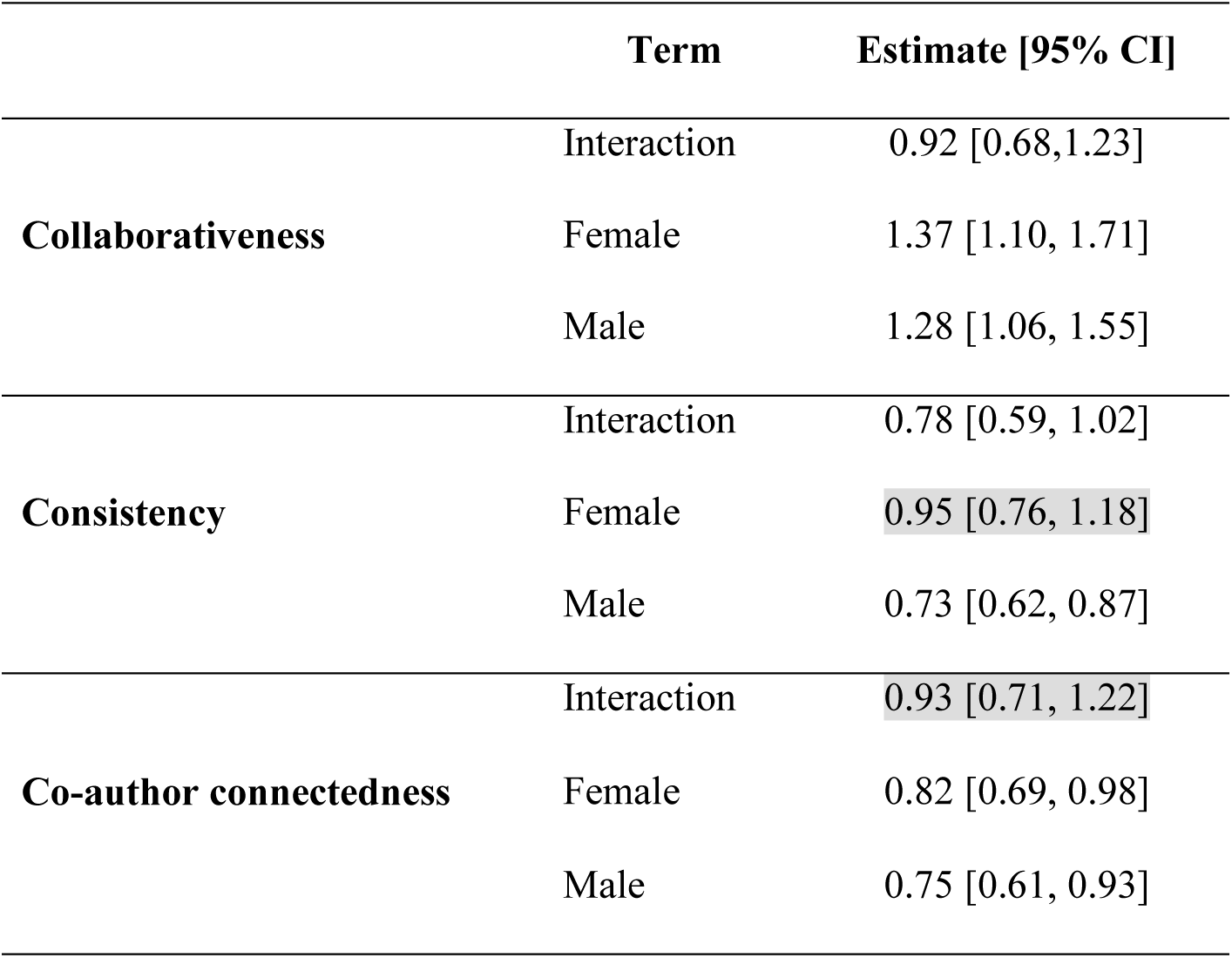
Output from AFT models testing how career length is influenced by the three measures of co-authorship sociality, as calculated over the first ten years for authors that started publishing just before or after their first ISBE conference attendance. Shaded estimates indicate non-significant relationships (95% confidence intervals overlapping 1.0).

|  | Term | Estimate [95% CI] |
| --- | --- | --- |
| **Collaborativeness** | Interaction | 0.92 [0.68,1.23] |
|  | Female | 1.37 [1.10, 1.71] |
|  | Male | 1.28 [1.06, 1.55] |
| **Consistency** | Interaction | 0.78 [0.59, 1.02] |
|  | Female | 0.95 [0.76, 1.18] |
|  | Male | 0.73 [0.62, 0.87] |
| **Co-author connectedness** | Interaction | 0.93 [0.71, 1.22] |
|  | Female | 0.82 [0.69, 0.98] |
|  | Male | 0.75 [0.61, 0.93] |

## Notes

http://www.tiny.cc/vanderwal

